# A layer cake model for plant and metazoan chromatin

**DOI:** 10.1101/2023.02.24.529851

**Authors:** Léoplod Carron, Lorenzo Concia, Stefan Grob, Fredy Barneche, Alessandra Carbone, Julien Mozziconacci

## Abstract

Early structural studies of chromosome organisation in eukaryotic nuclei led to the identification of euchromatin and heterochromatin, two main archetypes of chromatin visible through microscopy. Subsequently, the diversity of chromatin composition along the linear genome sequence was resolved using high throughput sequencing techniques to map a multitude of chromatin marks and unravel their functional organisation. Recent analyses of human chromosome conformation capture experiments tend to point to the existence of several chromosome sub-compartments that may correspond to epigenomic variations in chromatin composition. Here, we compare genome 3D organisations in representative eukaryotic species to explore the links between chromosomal sub-compartments and chromatin marks, genome replication timing, and genomic repeats in six model organisms, including vertebrates, plants and insects. We report that the 3D organisation of chromatin in organisms with different genome content and size can be described as layers characterised by distinct chromatin marks and activities. We propose a ”layer cake” model for the genome 3D organisation as a more refined view than the prevalent ”two compartments” model of chromatin organisation in multi-cellular organisms.

## Introduction

Chromatin is formed by the association of DNA and histones in the nucleus. The name chromatin was originally coined by Walther Flemming in the 19th century and it constitutes today the subject of a very active field of research due to the role of chromatin in genome packaging, gene expression and cell epigenomic inheritance. Electron microscopy observations of the interphase nucleus led to the first distinction between heterochromatin, which comprises the dense regions of chromatin, and euchromatin, often located in the centre of the nucleus and less dense. These two types of chromatin were identified as respectively repressing and permissive to transcription and are since then recognised as key players to sustain cell-type-specific genome 3D conformation and transcription. With the rise of high throughput sequencing, the mapping of the epigenome revealed a mechanistic link between chromatin functional status and its composition and organisation. Among characteristic histones and histone post-translational modification, histone H3 di- or trimethylation (H3K9me2/H3K9me3) are enriched at condensed and silent constitutive heterochromatin whereas histone acetylation is typically found at transcribing genes in euchromatin. The case of H3K27me3, which co-localize with the silencing factor Polycomb, is somewhat subtle since it is found in smaller islands within euchromatic active marks. These regions have been termed “facultative heterochromatin” because they can be either expressed or repressed in different cell types compared to “constitutive heterochromatin” regions which are usually considered as “constitutively inactive”.

Over the past decade, Chromosome Conformation Capture (3C) experiments, which map the contact frequencies between different loci along chromosomes, have linked the classical euchromatin/heterochromatin view obtained from microscopy experiments to the epigenomic maps. The main result of the first, and low resolution, genome-wide 3C analysis (Hi-C) of intact human nuclei was that euchromatin and heterochromatin were forming two spatially distinct compartments, referred to as A and B, which strongly correlate with gene expression [16]. Interestingly, facultative heterochromatin, which contains repressed genes, was at the time found together with euchromatin in the A compartment. This organisation in two compartments is conserved in multi-cellular eukaryotes, including plants [8] and species with holocentric chromosomes [7]. Principal features of chromatin compartmentalisation were later on shown to correlate with replication timing[24], TE content [5] and many other epigenetic marks [26].

Improvements in Hi-C resolution have recently allowed to shed new light on nucleus 3D organisation and its link to chromatin marks, notably suggesting existence of intermediate sub-compartments within A and B compartments (that we will call here I for “intermediate”) [24, 28, 17, 21, 3]. Interestingly, these studies point towards a strong enrichment of facultative heterochromatin in the I compartments. The number of sub-compartments considered in these studies ranges from 3 to 6. Here we compare and contrast chromatin compartments in five distantly-related multi-cellular species based on Hi-C data. We correlate compartments with distinctive histone marks and DNA repeat families and report a common organising principle of 3D chromosome architecture as a multi-layered compartment structure. These layers, corresponding to the formerly described compartments ranging from A to B through I, come with different chromatin flavours and are conserved in all species studied, with a main difference between relatively short, more compact genomes, and longer genomes. In the plant *Arabidopsis thaliana* and the fly *Drosophila melanogaster*, for which estimated genome sizes are 135 and 180 Mbp long, respectively, TEs are enriched in peri-centromeric regions and are more dispersed in the chromosomal arms which exhibit a compartment layer-like organisation. In human, mouse and chicken, for which the estimated genome sizes are in between 1 and 3 Gbp, TEs are found scattered throughout the whole genome, in the A, B and I compartments, with specific families being enriched in A and B compartment types [5].

## Results and discussion

### Layered compartment organisation of human chromatin

In this study, we performed a detailed analysis of chromatin compartments in five representative species of metazoans and plants and highlight the existence of A/B intermediary states that subtly partitions epigenomic features. We hereafter start by illustrating this partition in human chromatin and later extend it to Mouse, Drosophilia and Arabidopsis Thaliana. Two emblematic features can be observed on the correlation matrix obtained from one (here the 16th) of the intra-chromosomal Hi-C contact maps from human nuclei (Fig. 1A). The first one is the white cross in the middle, which corresponds to repetitive, un-mappable (peri-) centromeric regions which are filtered out from further analysis. The second one, which was already reported in the founding Hi-C study [16], is the existence of a checker-board pattern resulting from alternating regions of two different types exhibiting a strong affinity for the same type. This pattern is interpreted as an indication of the presence of two spatially disjointed compartments, which correspond to eu- and heterochromatin hallmarks. In line with this initial study, and with the many to come, we computed the first eigenvector of the correlation matrix (Fig. 1B). When properly oriented, a positive value of the eigen-vector corresponds to the A, euchromatic compartment and a negative value to the B, hetero-chromatic compartments (respectively in red and blue on Fig. 1B). When reordering the matrix according to this eigenvector values, the resulting map shows that B and A regions, respectively at the top left and bottom right, have a highly correlated contact profiles, while the contact profile between regions from the B type and the A type are negatively correlated. Since the transition between A and B is progressive, we identify additional compartments within A, B and the A/B transition zone. To this end, we used a six states Hidden Markov Model (HMM) trained on the original correlation map. The resulting states were found to alternate along the chromosomal arms (Fig 1D). When ordered with respect to the first eigenvector, we can see that these states correspond to gradual transitions in the transition region from A to B (Fig 1E). The number six that we chose is here arbitrary and could be higher or lower, the only requirement being that this number should be high enough to describe the gradual change of the first eigenvector values when going from A to B compartments.

**Figure 1:**
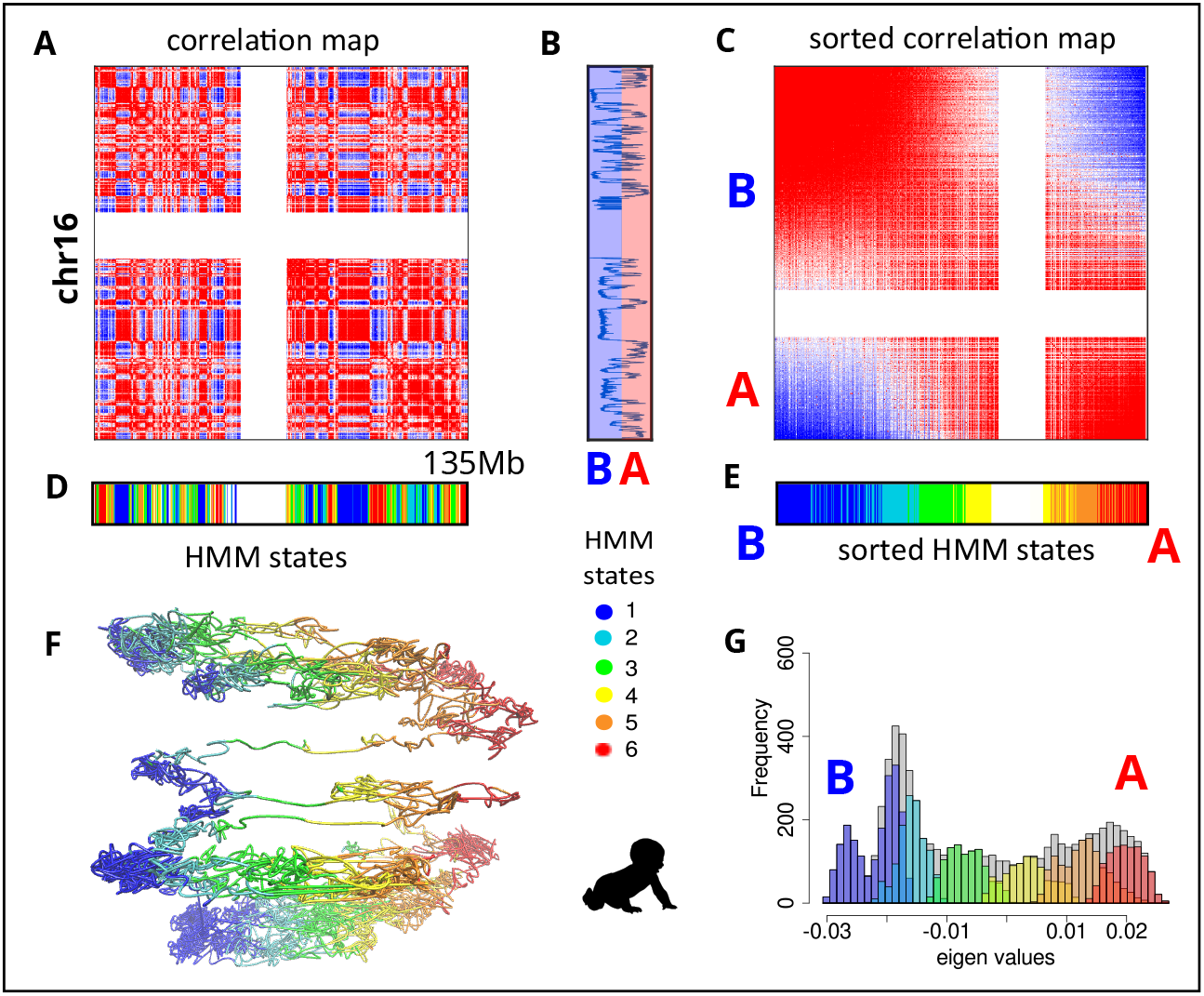
3D chromosome structure exhibits a layer cake pattern **A** Pearson correlation map *C* of human chr 16 at a resolution of 100kb. **B** First eigenvector of *C*. **C** Pearson correlation map *C* sorted by the first eigenvector. **D** The six states obtained with the HMM. They are coloured blue, light blue, green, yellow, orange and red. **E** The six states found in D sorted by the first eigenvector, as in C. **F** Average 3D structure of chromosome 16 coloured with the six states obtained with the HMM, as in D. **G** Distribution of the values of the first eigenvector (grey). The distribution for each of the six states is shown with the colours corresponding to each compartments. Color-scale as in D, centromeres have been filtered out.

In order to illustrate how the states are organised in 3D, we coloured a 3D representation of the contact map obtained with ShRec3D [15, 19]. Since Hi-C data come from a population of cells, this 3D structure represents an average consensus representation of the data. The six HMM states form layers in 3D, stacked from the most B-like to the most A-like (Fig 1F). We use here the term layer that differs from the term compartment since the genomic regions in these layers are not completely isolated from each others, but rather gradually organised in the 3D space. Reflecting the observed 3D arrangement of the regions in layers corresponding to the different states, regions that are found in the same state also exhibit similar values of the first eigenvector. The distribution of these values for each state illustrates the gradual changes from B to A (Fig1G). The classical distinction between region from the A and B compartment is done by looking at the sign of the value of the first eigenvector, the distribution in Fig1G is nevertheless far from bimodal, as would be expected in presence of two compartments, but rather in agreement with a gradual layered organisation as described in this paper, with layers 1,2,3 and 4,5,6 respectively corresponding to B and A [17].

In order to tackle the question of the spatial organisation of these layers in the nuclear space we used the results from the SPIN methodology [31], an integrative computational method to reveal genome-wide intra-nuclear positioning of genomic loci relative to nuclear speckles, lamina, and putative associations with nucleoli. We recovered that the six layers strongly correlates with SPIN states, although these were obtained on a different human cell type (K562, Fig S1). The most B-like 3D layer (state 1 of our HMM model) corresponds to the lamina-associated SPIN state, whereas most A-like 3D layer (state 6 of our HMM model) is associated with the SPIN comprising nuclear speckles and active chromatin regions at the interior of the nucleus. Intermediate layers 2-3 are associated with lamina and near lamina states while layers 4-5 are associated with euchromatic interior states (Fig S1). This comparison shows that while they are only derived from Hi-C contacts, HMM-defined states closely mirror SPIN states, which are derived from a larger set of experimental data. This similarity prompted us to investigate the organisation of chromatin in other species and assess the conservation of the layered organisation described here in human cell lines.

### Layered compartment organisation in plants and metazoans

In order to know whether the organisation of chromosomes is similar in other metazoans and plants we applied the same methodology on Hi-C data from mouse, chicken, drosophila and *A. thaliana* (Supp Fig 1, Fig 2). Our results showed the presence of a checker-board pattern in the correlation maps (Fig 2, bottom) as well as a layered organisation of HMM-derived compartments on the average 3D structures (Fig 2, top). In all species studied here, the genome is alternating between two ground states 1 and 6, hence the checker-board pattern, with a smooth transition between those two states, hence the layered organisation.

**Figure 2:**
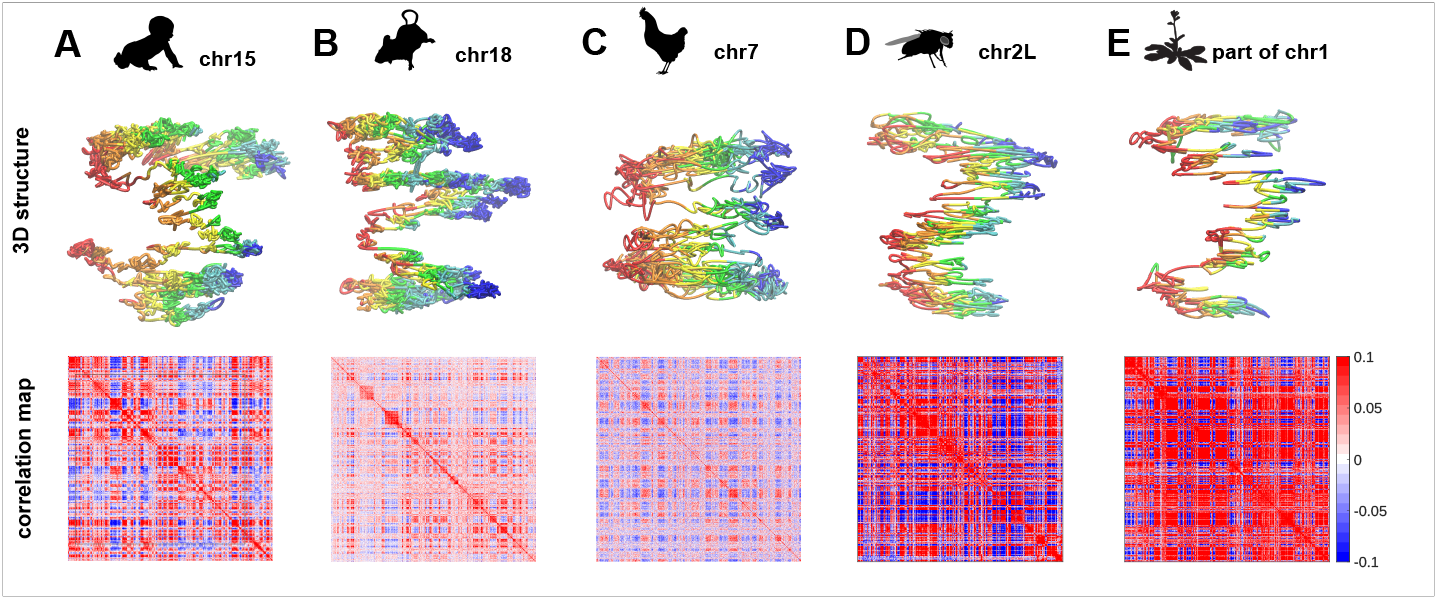
Universality of the layer cake organisation in metazoan and plant chromosomes Overview of averaged consensus 3D structures from different species at 10kb resolution with the overlap of the six HMM-derived states. For each chromosome or segment of chromosome, the figure includes the 3D representation labelled with 6 layers (top) and the correlation map (bottom). **A** Human chromosome 15. **B** Mouse chromosome 18. **C** Chicken chromosome 7. **D** Drosophila chromosome arm 2L. **E** A. thaliana chromosome 1 (region between 16 and 23 Mbp).

However, we observed a difference between the organisation of these compartments in longer genomes (human, mouse and chicken) compared to shorter ones (Drosophila and *A. thaliana*). While the checker-board pattern could be seen at large scale (i.e. on several chromosomes or on a full chromosome) for long genomes, it could be retrieved when zooming on shorter regions of the chromosomal arms for shorter ones (Fig S2). This specific feature of shorter genomes is not specifically due to the genome size but to the higher mappability of (peri-)centromeric sequences in the smaller genomes [22]. When they are included in the Hi-C maps, mappable centromeres form squares of high contact frequency values that are spatially dissociated from chromosomal arms [20]. The correlation matrix obtained from the Hi-C contact map thus reflects this specific organisation, as highlighted by the first eigenvector which delineates the position of the centromeres (Fig. S2, left). When the centromeres are excluded from the contact maps by zooming in on the chromosomal arms, the checker-board pattern characteristic of the presence of A/B compartments appears and correlates with gene activity as previously reported by Dong & al. [8] in *Arabidopsis Thaliana* (Fig. S2, right). In human, mouse and chicken, centromeres consist of highly repetitive and therefore masked sequences and it is not necessary to zoom on the chromosome arms to extract the checker-board pattern.

### Layered compartments correspond to different epigenomic features, DNA Repeat contents and replication timings

Now that we have split the genomes of these different species into seven regions, the 6 layers given by the HMM together with the regions that have been filtered out, corresponding mainly to the (peri-) centromeres of the chromosomes, we used publicly available epigenome profiling datasets to test whether these regions correspond to distinct chromatin states. For this purpose, we first used the chromatin states that have been previously described for Human [9], Drosophila [10] and *A. thaliana* [25] chromatin. These states were obtained using 1D annotations of the genomes, corresponding mostly to histone modifications and association with the lamina. For human (Fig 3A), we find that active chromatin marks are enriched in the A-like layer 6 while inactive chromatin is enriched in the B-like layer 1. We also see that the three states corresponding to polycomb repressed chromatin are enriched in intermediate layers 3 and 4. This finding is in agreement with recently published studies [2, 28, 18, 21] and suggests that polycomb and bivalent domains, that can be activated or inhibited depending on the cellular context, are typically located in between constitutively active and inactive regions. Interestingly, we found that this particular positioning of polycomb is conserved in Drosophila (Fig 3B) and *A. thaliana* (see also [12], Fig 3C), suggesting that this property could be a hallmark of all species that have polycomb. Of note, the tendencies observed in Fig 3 do not depend on the number of layers that was chosen but are rather due to the distribution of chromatin marks along the chromosomes.

**Figure 3:**
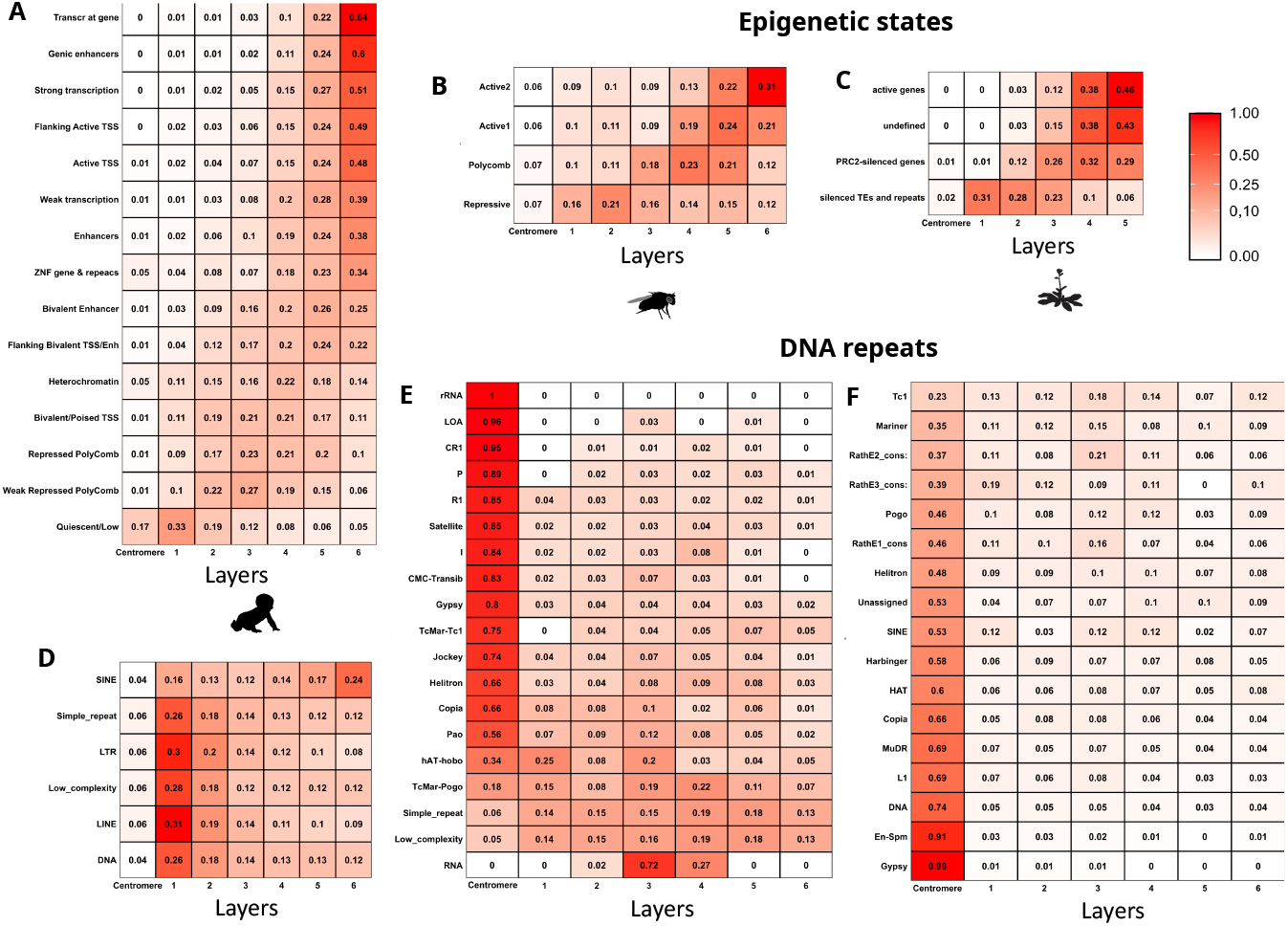
Chromatin and sequence composition of the layers. **A** Epigenomic annotations (ChrommHMM[9]) in Human. **B** The chromatin colors [10] in Drosophila. **C** Epigenomic annotations [25] marks in Arabidopsis Thaliana. **D**,**E**,**F** Enrichment in repetive sequences in the layers in Human, Drosophila and Arabidopsis [1]

We finally checked for the enrichment of DNA repeats families within the different layers described here. In mammalian chromosomes, the enrichment of SINEs in the A compartment and LINES in the B compartments [5] is reflected by a correlation between these DNA repeats and the layer organisation (Fig 3D). On the other hand, for shorter genomes such as the Drosophila and Arabidopsis, the majority of repeats (tandem repeats as well as dispersed repeats) are found in the (peri-)centromeric regions and we did not find any repeat family significantly over-represented in any of the layers (Fig 3FE). The few repeats families that were not found to be enriched in the (peri-)centromeric regions were preferentially found in the intermediate layers (2 to 5) in both Drosophila (e.g. low complexity and simple repeats) and Arabidopsis (e.g. Tc1 and Mariners).

Taken together, our results highlight the existence of chromosome architecture as 3D layers corresponding to different “flavours” of chromatin and DNA sequence. This partition goes beyond the mere distinction between eu- and hetero-chromatin, with the noteworthy observation that Polycomb repressed domains were on average found in intermediate layers between active and inactive chromatin. We also described here a difference between relatively shorter, more compact, genomes and longer genomes. In Arabidopsis and Drosophila, for which the genome is respectively 135 and 180 Mbp long, DNA repeats are preferentially found in peri-centromeric regions while in mammals, for which the genomes are 3 Gbp long, these elements are found scattered in the whole genome with specific families being enriched in the extreme layers 1 and 6. The origin of this difference remains an important question and the relationship uncovered here between the repeat content, the chromatin composition and the 3D genome folding adds a new perspective to this matter.

## Material and Methods

### Datasets used in the present study

All the datasets used here are summarised in figure Supplementary Table S1. We downloaded Hi-C from Human, Mouse, Chicken, Arabidopsis thaliana and Maize genome [14, 24, 11, 8, 23, 29] from SRA archive and reanalyse them with Hi-C pro[27]. For chromatin colors, we used the datasets presented in [9, 10, 25]. For the replication timing, we used the datasets presented in [13, 32, 4].

### Hi-C analysis

HiC data where downloaded from previously published study described in Supplementary Table S1. Hi-C reads were processed using HiC-Pro[27]. All generated map were produce at a resolution 10kb and 100kb.

To extract A and B compartments, we first normalise the intra-chromosomal contact count matrix by Sequential Component Normalization (SCN)[6] in order to have the same sum of contact in each row and column. Then each contact is divide by the mean number of contact of his diagonal[30] in order to obtain the matrix ratio of observed contact under expected. The first eigenvector of the correlation matrix *C* is then oriented with respect to the gene density to get the A and B compartments [16].

3D structures were produced with the Matlab (https://www.mathworks.com/products/matlab.html) implementation of ShRec3D[15] with an alpha parameter of 0.2 and using the “strain” dimension reduction method.

### Compartment extraction using the Hidden Markov Model

We extracted the genomic compartments with a hidden markov model (HMM) as described in Rao *et al*.[24] using the HMMlearn library in python direclty on the correlation matrix *C*. We used the gaussianHMM function with parameters *covariance=‘diag’, niter=100* using 6 component in the model. We then used the function *predict* in order to have the most probable compartment sequence. Since compartment states from HMM are not ordered we used the first eigenvector of the correlation map as computed in the previous section to order them from the most “A-like” to the most “B-like”. Fro comparison, we also generated compartments from inter-chromosomal contacts, using the same pipeline.

### Partition of DNA repeat families, epigenomic states and replication timing in the different regions of the genome

For each dataset (Table S1), we downloaded bed files in order to have the genome coverage of each tracks. We summed the signal over 10kb non-overlapping bins for each track and computed the overall signal in each compartment, as determined using HMM. This sum was then normalised to one in order to get the proportion of the feature of interest in each of the six compartments

## Acknowledgements

The authors are very grateful to Vincent Colot teams for precious advise during common lab meetings; Irc chat of bioinfo-fr canal for technical guidance with repeat element. Work by L Carron was funded by a collaborative research grant from Agence Nationale de la Recherche ANR-18-CE13-0004-01 to AC and FB. Work in FB laboratory was also supported by a research grant from the Velux Foundation (Switzerland), the CNRS EPIPLANT Action (France) and by the COST Action CA16212 INDEPTH (EU).

## Author’s contributions

LeC developed the Hi-C analysis tools, analysed Hi-C datasets, prepared annotation and made the figures. LeC and LoC analysed the Chip-seq experiments in Arabidopsis thaliana. FB, AC and JM supervised research. LeC, and JM wrote the manuscript with input from FB, LoC, AC and SG.

---

**Table S1:**
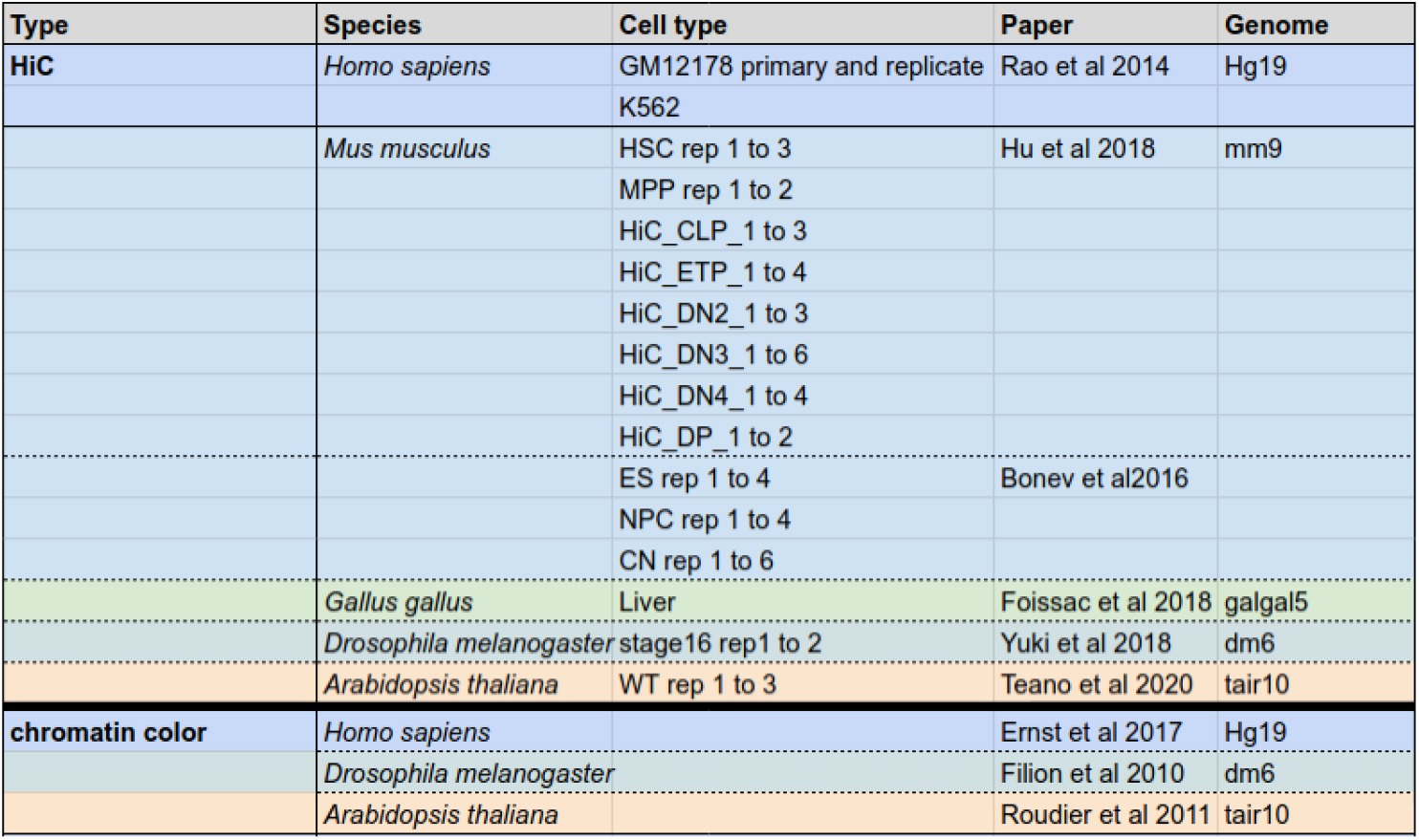
Table of Hi-C data-sets used this study with the reference genome used in the mapping step.

**Figure S1:**
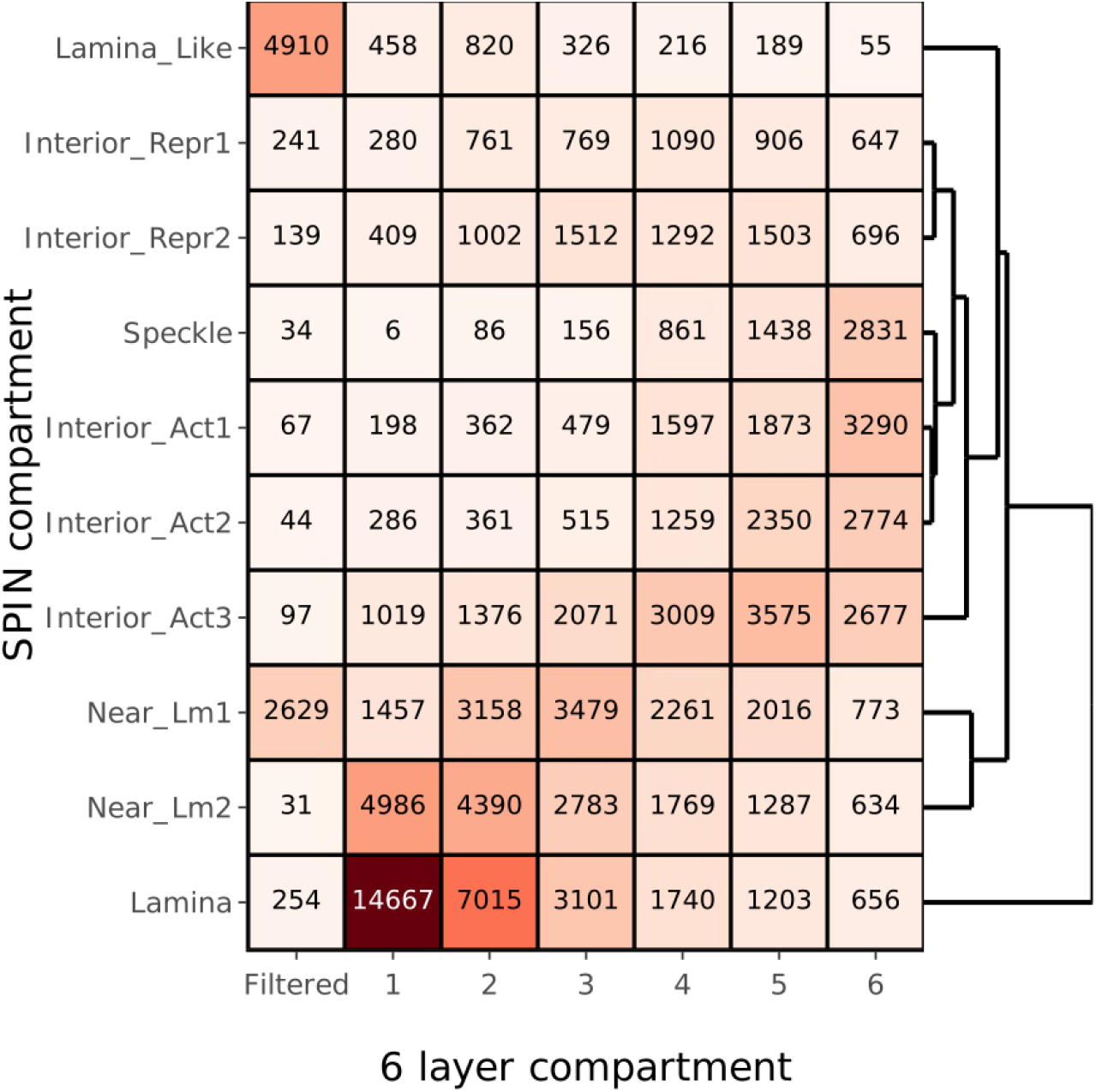
Heatmap of contingency between the SPIN compartments and the 6 layers inferred from the intra-chromosomal Hi-C contacts in human cells at a resolution of 25kb.

**Figure S2:**
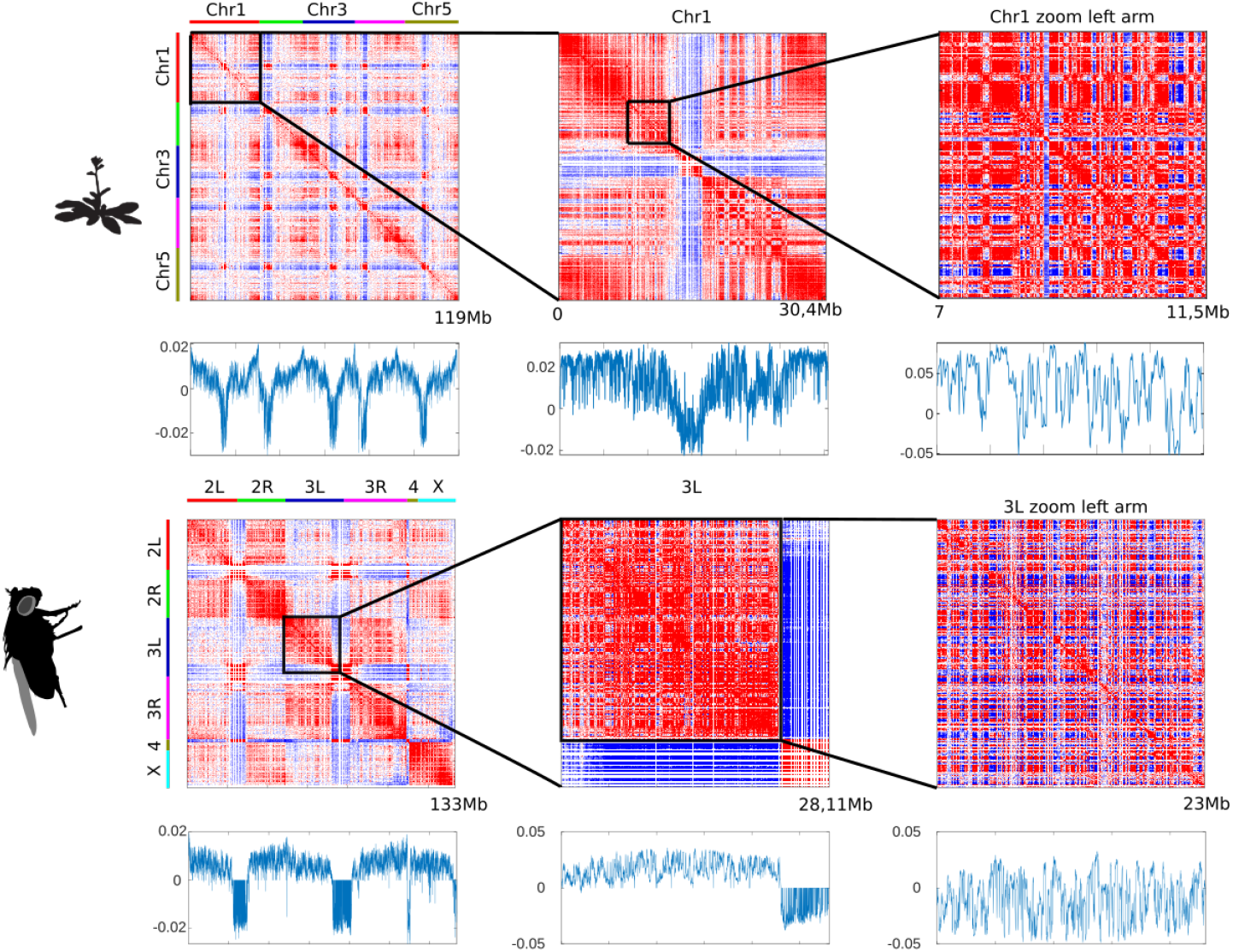
Hi-C Pearson correlation maps (top) for *Arabidopsis thaliana* and *Drosophila melanogaster* at a resolution of 10kb. Correlation matrices and the first eigenvectors (below each matrix) were computed at three different scales, from several chromosomes to zooms on specific regions of the chromosomal arms.

